# Sex-specific effects of chronic unpredictable stress on mitochondrial function in the HPA axis in mice

**DOI:** 10.1101/2025.07.28.667247

**Authors:** Alexia M. Crockett, Noelle I. Frambes, Alaina Mullaly, Amelia M. Churillo, Rinaldo R. Dos Passos, Cameron Folk, Lisa Freeburg, Eliana Cavalli, Josephine Gardiner, Evelynn N. Harrington, Timothy L. Philbeck, Stephanie Wilczynski, Fernanda Priviero, Francis G. Spinale, R. Clinton Webb, Susan K. Wood, Michael J. Ryan, Fiona Hollis

**Affiliations:** University of South Carolina School of Medicine, Department of Pharmacology, Physiology & Neuroscience; Columbia VA Health Care System; University of South Carolina School of Medicine, Department of Cell Biology and Anatomy; Department of Biomedical Engineering, Molinaroli College of Engineering and Computing, University of South Carolina; The Cardiovascular Translational Research Center, University of South Carolina; The Institute for Cardiovascular Disease Research, University of South Carolina

## Abstract

Stress, whether real or perceived, activates physiological and behavioral responses via the hypothalamic-pituitary-adrenal (HPA) axis and sympathetic nervous system activation. Under chronic stress, however, these adaptive responses become dysfunctional leading to pathological changes in behavior and health. Mitochondria are dynamic organelles essential for cellular energy production and for initiating glucocorticoid synthesis and release from adrenal glands during stress. Thus, mitochondria may represent a first line of response to environmental challenges. However, the effects of chronic stress on mitochondrial function within the HPA axis, particularly regarding sex differences, are unexplored. We exposed adult male and female C57BL6/J mice to four weeks of chronic unpredictable stress and examined behavioral and mitochondrial responses in the hypothalamus and adrenal glands – two key HPA axis regions. As previous reports indicated sex differences in stress responsivity, we hypothesized that chronic stress would differentially impact mitochondrial respiration within HPA axis regions in a sex-specific manner. Chronic stress increased avoidance behavior in males and passive coping behavior in females, indicating sex-specific behavioral responses. In females, stress significantly decreased mitochondrial respiration in both the hypothalamus and adrenal glands, while males were not significantly affected. In males, stress increased adrenal expression of mitochondrial complex II protein, which may have served a compensatory role to preserve mitochondrial function. Mitochondrial respiration significantly correlated with behavioral measures in stressed animals, highlighting a relationship between metabolism and stress-induced impairments. These findings reveal sex-specific metabolic adaptations to chronic stress and suggest that females may be more vulnerable to stress-induced mitochondrial dysfunction within the HPA axis.

**Clinical Perspectives:**

- Chronic stress is widely prevalent, associated with neuropsychiatric disease that affect women at a rate twice as high as men, and mediated by mitochondria, yet sex differences in the effects of chronic stress on mitochondrial function have not been characterized.
- Despite similar behavioral outcomes, chronic unpredictable stress exposure significantly decreases mitochondrial respiration only in the hypothalamus and adrenal glands from females, in association with stress-induced behavioral alterations.
- Females may have increased vulnerability to metabolic effects of chronic stress and therapies targeting mitochondrial function may be more efficacious in preventing behavioral impacts of stress in females.

## 1. Introduction

The population of the United States is increasingly under chronic stress, with 1 in 4 adults reporting that stress levels impede their daily functioning (1). Chronic stress is linked to the onset and severity of various pathologies, including psychiatric disorders (2) like depression and anxiety, resulting in an estimated economic burden of around $280 billion (3). During a stressful or threatening event, the body tries to regulate physiological processes to conserve resources as protection. These actions are mediated by the hypothalamic-pituitary-adrenal (HPA) axis, a coordinated neurobiological response system that triggers an increase in circulating glucocorticoids, leading to physiological and behavioral adaptations as well as the generation and reallocation of energy resources to facilitate a return to homeostasis (4,5). Although the HPA axis is under negative feedback regulation (4), chronic stress exposure can induce dysregulation (6), leading to maladaptive behavioral and molecular responses. Moreover, the responses of the HPA axis can be context-, time-, and sex-specific (7). Women report higher levels of stress and are twice as likely to suffer from stress-related psychiatric disorders compared to men (8,9), yet female subjects are underrepresented in stress research and the mechanisms underlying these sex differences are poorly understood.

While mitochondria are commonly understood for their central role in energy production in the form of ATP, they are also crucial for processes related to cell survival and health including biosynthesis of steroid hormones (10,11), production and regulation of reactive oxygen species (ROS) (12), and apoptosis (13). Mitochondria also release immunomodulatory molecules such as mitochondrial DNA (mtDNA) and ROS (12), resulting in inflammasome activation and subsequent pro- and anti-inflammatory cytokine release. In addition to housing the rate-limiting enzymes for corticosteroid synthesis, mitochondria are also sites of action of steroid hormones, as glucocorticoids have been found to directly interact with the mitochondrial genome (14,15) and influence mitochondrial function in neurons (7). The impact of glucocorticoids on the brain are especially crucial given that the brain utilizes 20% of the body’s basal oxygen consumption (16), making it highly sensitive to alterations in mitochondrial function, specifically during chronic stress, when energy requirements are increased (17). Moreover, we previously demonstrated that small fluctuations in brain mitochondrial function can drive behavior (18–20). A disconnect between energy demand and mitochondrial respiratory function during energy intensive experiences, such as chronic stress exposure, could have significant consequences leading to pathology, yet the relationship between chronic stress, brain mitochondrial function, and stress-induced behavioral pathology is unclear. The chronic unpredictable stress (CUS) paradigm is a well validated model used to assess the effects of chronic stress on behavioral pathology and physiology in rodents (21–24). CUS has been shown to alter brain function, including metabolism (reviewed in (25)). Preclinical studies have found significant effects of chronic stress exposure on mitochondrial function across several brain regions in males (26–33). However, the exclusion of females from these studies has led to a scarcity of information regarding sex differences in the response of mitochondria to chronic stress.

We aimed to address these knowledge gaps by analyzing the effects of chronic stress exposure on mitochondrial function within the hypothalamus and adrenal glands, as these represent the entry and exit points of the HPA axis. We hypothesized that CUS exposure would induce sex-specific disruptions to mitochondrial function in HPA axis regions, resulting in behavioral pathology and peripheral dysregulation. To test this, we exposed male and female mice to 28 days of CUS. We then evaluated the impact of the CUS exposure on avoidance and stress-coping behaviors, mitochondrial respiration and complex protein expression in HPA axis regions, and endocrine release. Our results show distinct sex-specific disruptions to behavior and mitochondrial function within the hypothalamus and adrenal glands, and significant correlations between these measures. These results highlight a role for mitochondria in mediating stress-induced alterations in the brain and behavior.

## 2. Materials and Methods

### 2.1 Animals

Adult (8 weeks old) male and female C57Bl/6J mice were used. All animals were maintained on a 12:12h light cycle and had access to food and water *ad libitum*. Animals were group housed upon arrival to the animal vivarium and left to habituate for one week and then handled for three days prior to the start of stress. Once the stress paradigm began, animals were randomly separated into control (CON) or stress (CUS) groups. CON animals maintained group housing, while CUS animals were single housed. All behavioral experiments were conducted during the light phase between 0900h and 1230h. This study is part of a multi-laboratory collaborative project focusing on the impact of chronic stress on the brain and behavior (current study), the kidneys (co-author M.J. Ryan), and cardiac function (co-author F.G. Spinale). We used mice for measures across multiple studies to be consistent with humane experimental principles to reduce the number of animals involved in experimental research. Therefore, data for renal and cardiovascular assessment were collected in all animals but are not included in this manuscript as they will be published separately. Renal and cardiovascular assessments are described in the methods below and shown in Figure 1 to enhance reproducibility. All procedures were conducted in conformity with the University of South Carolina Institutional Animal Care and Use Committee and conformed to the U.S. National Institutes Health Guide for the Care and Use of Laboratory Animals.

**Figure 1.**
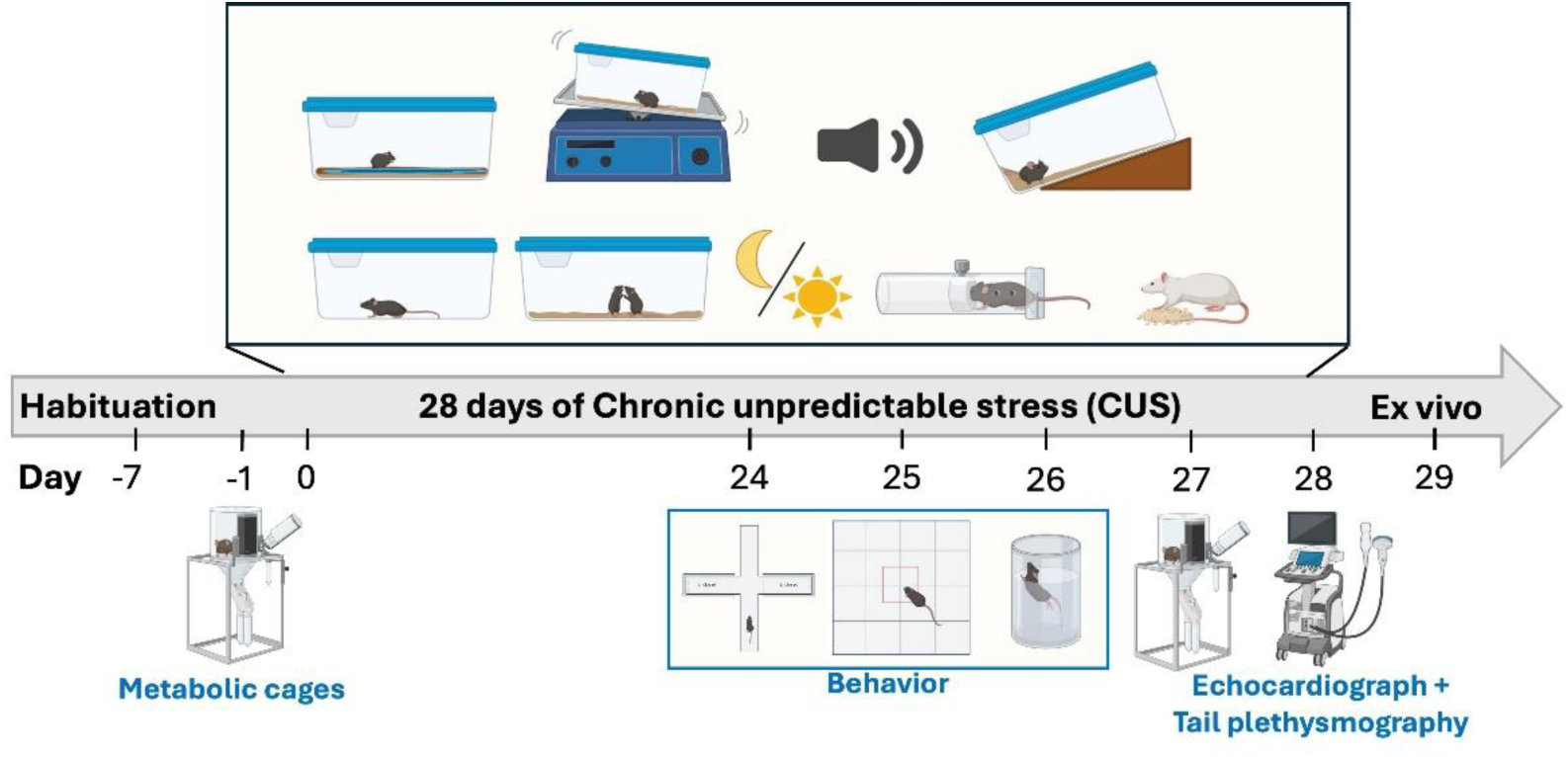
Experimental timeline. Following arrival, animals were allowed to habituate to the vivarium and experimenters with mild handling daily for one week (Days -7 – 0) and were weighed weekly to monitor changes in body weight. Animals were placed in metabolic cages for overnight urine assessment, one day prior to stress (Day -1) and one day prior to the end of stress (Day 27). Animals were then randomly assigned to chronic unpredictable stress (CUS) on control (CON) groups prior to the start of stress (Day 0). The Elevated Plus Maze (EPM) was measured on Day 24, the Open Field Test (OFT) was measured on Day 25, and the Forced Swim Test (FST) was measured on Day 26. On Day 28, animals were anesthetized, and blood pressure and cardiac function were measured via tail cuff and echocardiograph. On Day 29, animals were weighed and euthanized via rapid decapitation for trunk blood and brain collection for ex vivo analyses.

### 2.2 Chronic Unpredictable Stress Protocol

Male and female mice were exposed to chronic unpredictable stress paradigm (adapted from (21) for 28 consecutive days. The CUS protocol consists of both physical and psychosocial stressors under home cage isolation. The time and length of exposure of each stressor was varied to maximize unpredictability and uncontrollability, with no repetition of stressors within a given week. During stress exposure, animals were carefully monitored for signs of distress. **Table 1** indicates the schedule of stressors. Non-stressed control animals (“CON”) were handled weekly by experimenters. All animals were weighed weekly and monitored daily.

**Table 1.** Chronic Unpredictable Stress Protocol distribution across weeks with durations.

| Week | Stressors | Duration |
| --- | --- | --- |
| 1 | Cage tilt, overnight light, restraint, social stress, wet bedding, white noise, reverse L-D cycle | 8h, 11h30, 6h, 3h, 7h, 8h, 24h |
| 2 | Cage shake, soiled rat cage exposure, cage tilt, restraint, no bedding, white noise, social stress | 1h, 4h, 3h, 3h, 8h, 12h, 3h, |
| 3 | Restraint, overnight light, cage shake, social stress, wet bedding, soiled rat cage exposure, cage tilt | 4h, 11h30, 1h, 3h, 8h, 2h, 6h |
| 4 | No bedding, reverse L-D cycle, day of darkness, white noise, cage shake, restraint, social stress | 8h, 25h, 24h, 12h, 1h, 6h, 3h |

### 2.3 Behavioral Analyses

For each behavioral test, mice were habituated to the testing room at least 30 minutes prior to behavior.

#### 2.3.1 Elevated Plus Maze (EPM)

On day 24, all mice were tested for avoidance behavior in the Elevated Plus Maze. The testing arena was elevated (50 cm) from the floor and consists of two enclosed arms (35 x 5 x 40 cm) and two open arms (35 x 5 x 1 cm) which are separated by a center zone (5 x 5 cm). Lighting was maintained at 17-18 lux on the open arms. Mice were placed in the center facing a closed arm and allowed to explore all arms for 5 minutes. Animals were tracked using the automated Ethovision system (Noldus; Leesburg, VA, USA).). Center-, nose-, and tail- point were used to track animal’s movements. Avoidance behavior was determined by the percentage of time spent in the open arms of the platform over total time in the arena. Any animals that fell from the arena were excluded from analysis. The testing apparatus was cleaned with 10% EtOH solution before and after each animal.

#### 2.3.2 Open Field Test (OFT)

On day 25, all mice were tested for avoidance behavior and locomotor activity in the Open Field Test. The testing arena consisted of a large square arena (45 x 45 cm) with tall walls (40 cm) so that the animals could not escape. Lighting was maintained at 10-12 lux in the center of the arena. The arena was divided into virtual zones denoting the center and peripheral areas. Mice were allowed to freely explore the arena for 10 minutes. Animals were tracked using the automated Ethovision system (Noldus). Center-, nose-, and tail-point were used to track animal’s movements. Avoidance behavior was determined by the percentage of time the animals spent in the center over total time in the arena. The arena was cleaned with 10% EtOH solution before and after each animal.

#### 2.3.3 Forced Swim Test (FST)

On day 26, all mice were tested for stress coping style in the Forced Swim Test. Mice were placed into a large, clear Plexiglass cylindrical tank (20 cm in diameter, 30 cm in height) filled with water (26-28°C) up to 15 cm so that animals could not touch the bottom, nor escape. Animals were video-recorded for 6 minutes, and behavior was manually quantified at a later timepoint by an experimenter blind to the groups. Analysis included latency to become immobile and total time spent immobile. Latency to immobility included the first time at which the mouse ceased all movement relating to escape outside of minimal movement to continue floating. Immobility included the time in which the mouse remained in a stationary posture for more than 3 seconds, with minimal movements only associated with continuing floating.

#### 2.3.4 Behavioral z-score

To have an integrated measurement of behavioral phenotype, we calculated a z-score for test parameters measuring emotionality, as previously described (34–36). Individual z-scores were calculated for each test using the following formula:

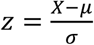

X represents the individual data for the observed parameter. μ and σ represent the mean and standard deviation for the control group, respectively. Here, as we were interested in the interaction between stress and sex, individual z-scores were calculated within each sex and normalized to same-sex controls. The time in the open arms for the EPM, time in the center for the OFT, latency to immobility in the FST, and body weight gain were included as individual test parameters for each mouse. Individual z-normalized raw scores were then averaged to generate a composite z-score for each sex. Within each of the test parameters, lower values indicate disrupted behavior. Thus, decreased or negative z-scores indicate worse emotionality or more stress-induced disruptions to behavior.

### 2.4 Renal Assessment

Overnight urine samples were collected from mice using metabolic cages prior to the start of the CUS protocol (day -1) and at the end of the protocol on day 27. Mice were removed from metabolic cages and returned to their home cages. Data from these assessments are in preparation for separate publication.

### 2.5 Cardiovascular Assessment

Heart geometry and function was assessed using a high-fidelity transthoracic echocardiogram (Vevo3100) on day 28, 24h prior to euthanasia. Animals were lightly anesthetized with isoflurane and placed on a heating pad during measurement. Shortly thereafter, systolic blood pressure was recorded in a subset of mice at the end of the study using tail plethysmography (Kent Scientific) under light isoflurane anesthesia. Mice were placed on an infrared heating pad to raise their body temperature to 37.5°C and tail cuffs were positioned to read for 5 cycles. Once animals regained consciousness, they were returned to their home cages. Data from these assessments are in preparation for separate publication.

### 2.6 Euthanasia and Tissue Collection

On day 29, animals were euthanized under basal conditions by cervical dislocation and rapid decapitation between 9:00-11:00am. Trunk blood was collected in EDTA-coated tubes (Sarstedt) and stored on ice. Plasma was collected from blood following centrifugation at 3,200 rpm for 15 minutes at 4°C. Plasma was snap frozen in chilled isopentane and stored at -80°C for later processing. The brain was rapidly removed, and both hemispheres of the hypothalamus and the adrenal glands were dissected out and weighed. Dissected sections were placed in a chilled well plate with 2mL of relaxing solution (BIOPS: 2.8 mM Ca2K2EGTA, 7.2 MK2EGTA, 5.8 mM ATP, 6.6 mM MgCl2, 20 mM taurine, 15 mM sodium phosphocreatine, 20 mM imidazole, 0.5 mM dithiothreitol and 50 mM MES, pH = 7.1) until later processing.

### 2.7 Mitochondrial Respirometry

Tissue samples were gently homogenized in ice-cold respirometry medium (MiR05: 0.5 mM EGTA, 3mM MgCl2, 60 mM potassium lactobionate, 20 mM taurine, 10 mM KH2PO4, 20 mM HEPES, 110 mM sucrose and 0.1% (w/v) BSA, pH=7.1) with a motorized Teflon pestle. 2 mg of tissue were used to measure mitochondrial respiration rates at 37°C using high resolution respirometry (Oroboros Oxygraph 2K, Oroboros Instruments, Innsbruck, Austria). A multi-substrate protocol was used to sequentially explore the various components of mitochondrial respiratory capacity as previously described (Hollis et al., 2015; Gorman-Sandler et al., 2023).

To measure respiration due to oxidative phosphorylation, various substrates were used to activate specific complexes of the electron transport chain. Coupled respiration through complex I was observed via addition of ADP (5mM) to a solution containing malate (2mM), pyruvate (10mM), and glutamate (20mM). Coupled respiration through complex II (CI+CII) was observed via addition of succinate (10mM). Electron transport system (ETS) maximal capacity was evaluated using the protonophore, carbonyl cyanide m-chlorophenyl hydrazone (CCCP; two successive titrations of 0.5 μM) until maximal respiration rates were reached. We then observed oxygen consumption in the uncoupled state by inhibiting complex I with the addition of rotenone (0.1 μM; ETS CII) to solely observe activity of complex II. Lastly, we inhibited electron transport through complex III with the addition of antimycin (2mM) to observe residual oxygen levels (ROX) due to oxidating side reactions outside of mitochondrial respiration. The O_2_ flux obtained in each step of the protocol was corrected for residual oxygen consumption (ROX).

### 2.8 Protein Expression

Relative protein expression was analyzed using Western blots, with primary antibodies (AB) against individual subunits of each complex in the oxidative phosphorylation (OXPHOS) system, Translocase of the Outer Membrane (TOM20) and 4-hydroxynoneal (4-HNE), a marker of oxidative stress. Protein concentration of hypothalamic and adrenal homogenates in respirometry medium was determined by BCA assay (ThermoScientific, Cat. #23227). Samples were prepared with 2X Laemelli buffer and milliQ water to a final concentration of 1μg/μL, then heated in a dry bath for 5 minutes at 37°C (for OXPHOS AB detection) or 95°C (for TOM20 and 4-HNE AB detection). 20μL of protein lysate was then loaded into 4–20% Criterion™ TGX Stain- Free™ precast gels (Bio-Rad, Cat. #5678094). Proteins were separated at 200V for 45 minutes and then transferred to a low-fluorescence PVDF membrane at 25V for 30 minutes using the Trans-Blot Turbo Transfer System and Kit (Bio-Rad, Cat. #1704275). Gels were activated by UV excitation using the Bio-Rad GelDoc XR + System prior to transfer, and membranes were imaged for total protein on the same system following transfer according to manufacturer’s instructions. TGX Stain-Free gels enable visualization and quantification of total protein across all lanes for normalization, comparable to Coomassie staining. Total protein was determined by measuring the entire lane and summing all the proteins present to account for differences in tissue type. Membranes were blocked using EveryBlot blocking buffer (Bio-Rad, 500mL, Cat. #12010020) for 30 minutes. Membranes were then incubated on a shaker overnight at 4°C with one of the following primary antibodies: Rodent Total OXPHOS, (Mitosciences, ab110413, 1:250), TOM20 (Cell Signaling, 1:1000, Cat. #42406), or 4HNE (Millipore, 1:1000, Cat. #AB5605) and Blocking Buffer. Following washes in PBS-T (VWR, Cat. #76371- 736), membranes were incubated with IRDye® 800 CW Donkey-anti-Rabbit (LI-COR, #926-6807), IRDye® 680RD Goat-anti-Mouse (LI-COR, #926–68070), IRDye ® 800CW Donkey anti-Goat (LI-COR, #926–32214), or IRDye® 800CW Goat anti-Rabbit (LI-COR, #926–32211) IgG secondary antibodies at 1:20000 dilution for 1 hour on a shaker at room temperature. After PBS-T washes, bands were imaged on a LI-COR Odyssey CLX System Imager. Captured bands were quantified as 16-bit TIF images using ImageJ software (NIH). The 4-HNE antibody binds to HNE-modified protein adducts formed during lipid peroxidation. As such, this results in multiple bands within each lane, of which only the most prominent band was quantified (70 kDa). To account for variation in sample loading, quantified bands were normalized to total protein. Western blot results are presented as fold change relative to same-sex control averages to account for between-blot variability.

### 2.9 Corticosterone Assay

Trunk blood was collected in EDTA-coated tubes and plasma was separated from blood by centrifugation at 3200 x rpm for 15 minutes at 4°C. Plasma was then used to assess CORT concentration using a commercially available enzyme-linked immunosorbent assay (ELISA) kit (Enzo Life Sciences, Cortisol ELISA kit, ADI-900- 071). Samples were diluted to a final concentration of 1:40 and analyzed in duplicates. Absorbance was read at 405 and 450 nm.

### 2.10 Statistical Analyses

Sample sizes are indicated in the figure legends. Experiments were run in 2 independent cohorts of n=8/group. Both cohorts followed identical experimental timelines, however sample sizes within behavioral and respirometry measures vary due to technical reasons (4 adrenal samples and 7 hypothalamus samples) or statistically significant outliers (as determined by the Grubbs Outlier Test – 1 mouse was excluded from mitochondrial respirometry). One male spontaneously died prior to the end of the study and was removed from all analyses. Data were analyzed by two-way ANOVAs (with stress exposure and sex as between-subjects factors), followed by Šídák’s multiple comparisons or uncorrected Fisher’s LSD post hoc test where appropriate. Pearson correlations were performed for composite z-score and mitochondrial respiration. Figures are shown as original, non-transformed data. All data were analyzed using Prism version 10 (GraphPad software Inc., San Diego, CA). P-values are reported in the main text with significance considered at the p≤0.05 level. Additional statistical details can be found in **Supplementary Table 1** for main text figures and **Supplementary Table 2** for supplemental figures.

## 3. Results

### 3.1 Validation of HPA axis dysregulation via CUS protocol

We first sought to validate the physiological and behavioral impact of our 4-week CUS protocol (**Figure 1**), given the substantial variation in published protocols. Previous studies using similar CUS protocols in mice reported that 4 weeks of CUS significantly altered body weight gain, exploration in the Open Field, passive coping and avoidance behavior (37–39) but did not alter adrenal gland weight (21,40). Therefore, we examined the same endpoints to benchmark our protocol’s efficacy.

#### 3.1.1 CUS decreases bodyweight gain in males but not females and did not affect adrenal gland weight

Exposure to 4 weeks of CUS revealed a significant stress x sex interaction on body weight gain (p=0.01) (**Figure 2A-B**). Post hoc analysis revealed that CUS significantly decreased body weight gain in males (p=0.01) but had no impact on stressed females (p=0.85). Similar to prior reports (21,40), we did not observe significant effects of stress (p=0.52) or a stress x sex interaction (p=0.34) on adrenal gland weight (**Figure 2C**). However, we replicated previous findings (41) of a main effect of sex (p=0.003) particularly evident in our stressed animals where stressed females had heavier adrenal glands compared to stressed males (p=0.005).

**Figure 2.**
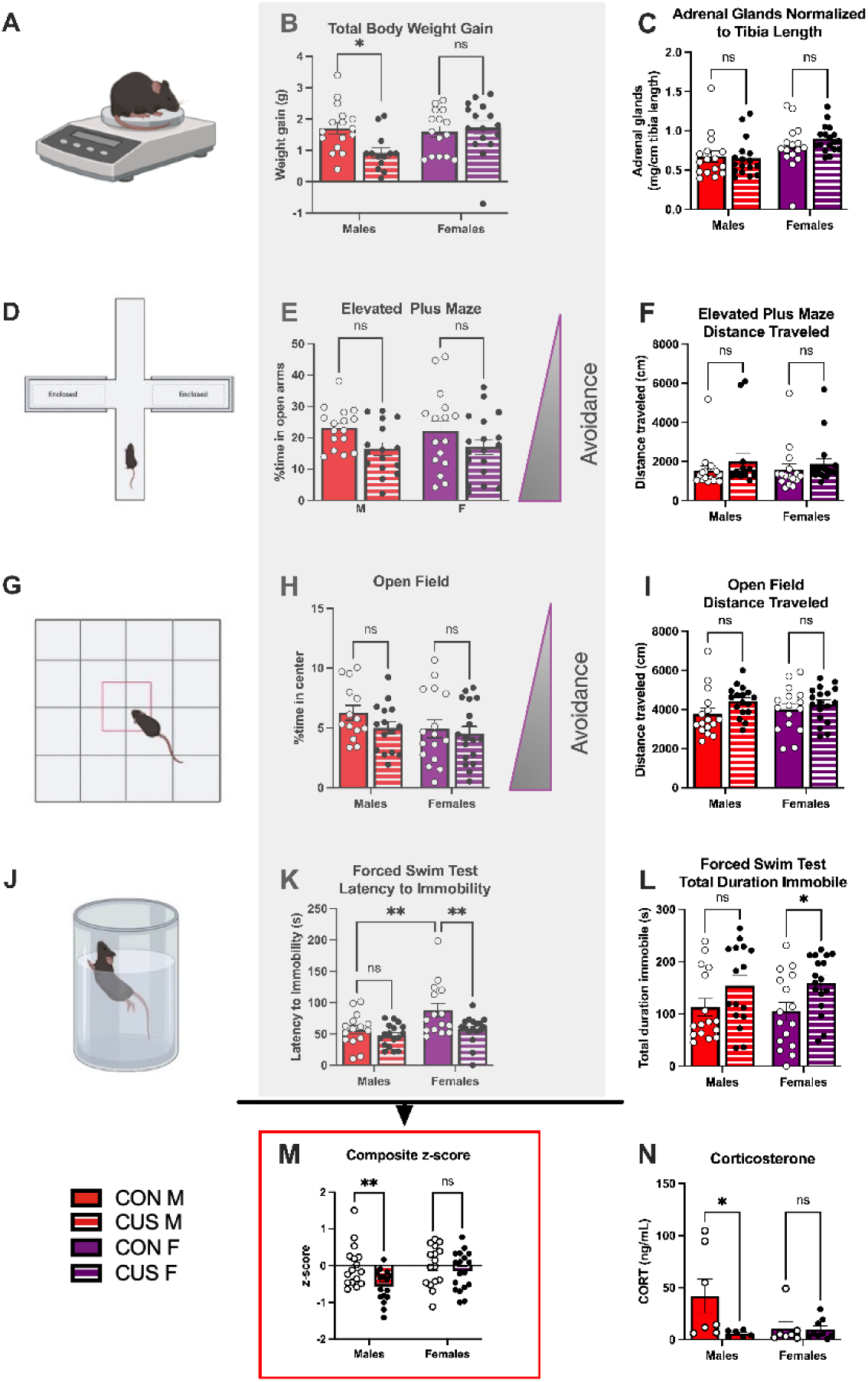
CUS induces significant behavioral and endocrine effects. **A)** Mice were weighed weekly and adrenal glands weighed post-mortem. **B)** CUS significantly decreased male total body weight gain, while females were unaffected. **C)** CUS did not significantly impact adrenal gland weight, however females overall exhibited heavier adrenal glands than males. **D)** In the Elevated Plus Maze (EPM), **E)** stress significantly decreased time spent in the open arms, while **F)** distance traveled was similar between CON and CUS groups. **G)** In the Open Field Test (OFT), **H)** CUS did not impact time spent in the center of the arena, nor **I)** the distance traveled. **J)** In the Forced Swim Test (FST), **K)** CUS significantly decreased latency to immobility, specifically in females. **L)** CUS increased the time spent immobile in both males and females. **M)** A composite z-score of body weight gain, EPM, OFT, and FST was calculated. The grey shaded region identifies the specific measures of each test that were included in the composite z-score. A lower score indicates more stress-induced behavior – decreased body weight gain, less time in the open arms, less time in the center, and decreased latency to immobility. CUS males had a significantly negative overall z-score compared to controls and there was no effect in females. **N)** CUS significantly blunted CORT levels, specifically in males. n= 16-18/group. Data are represented as ± SEM.

#### 3.1.2 CUS increases avoidance behavior and passive coping behavior in females

On day 24, we tested avoidance behavior in the Elevated Plus Maze (EPM; **Figure 2D**). We observed no main effect of sex (p=0.95) nor a stress x sex interaction (p=0.75). However, we did observe a main effect of stress to decrease the percentage of time spent in the open arms (p=0.01) (**Figure 2E**), distributed across both males and females. Notably, there were no significant effects of stress or sex on the distance traveled in the EPM (stress: p=0.21 and sex: p=0.92) and no stress x sex interaction (p=0.78) (**Figure 2F**). On day 25, we measured avoidance behavior and locomotor activity in the Open Field Test (OFT; **Figure 2G**). Contrary to others (21,37), we did not observe any significant effects of stress (p=0.19), sex (p=0.16), or a stress x sex interaction (p=0.50) on the percentage of time spent in the center (**Figure 2H**). Notably, we observed a trend for an effect of stress (p=0.07) to increase locomotion, but no effect of sex (p=0.83), or stress x sex interaction (p=0.41) on distance traveled in the OFT (**Figure 2I**). We assessed stress-coping behavior in the Forced Swim Test (FST; **Figure 2J**) on day 26, analyzing latency to and duration of immobility. Here, we saw a main effect of stress (p=.006) and sex (p=.005) to decrease the latency to immobility in CUS females (p=0.003) and a significant difference between CON males and females (p=0.003). However, we saw no stress x sex interaction (p=0.14) (**Figure 2K**). We also observed a main effect of stress to significantly increase the duration of time spent immobile (p=0.005) which was primarily driven by stressed females (p=0.04) (**Figure 2L**). We did not observe main effects of sex (p=0.95) or a stress x sex interaction (p=0.68) on time spent immobile.

#### 3.1.3 Stressed males but not females exhibit significant stress-related behavioral impairments

We generated individual z-scores for body weight gain and each behavioral test (**Supplemental Figures 1A-D**) and, when integrated together as a composite z-score, we observed a main effect of stress to decrease overall composite z-score (p=0.001) (**Figure 2M**). Since lower z-scores indicate greater expression of stress-related behavioral impairments, this finding reflects a negative impact of CUS on overall behavioral outcomes. Post hoc analysis revealed this effect was primarily driven by stressed males (p=0.01) as stressed females did not exhibit a significant difference from CON counterparts (p=0.16). We observed no main effects of sex (p=0.11) and no stress x sex interaction (p=0.38).

#### 3.1.4 CUS decreases basal plasma corticosterone levels in males but not females

We analyzed resting plasma corticosterone levels and found a main effect of stress to decrease plasma CORT levels (p=0.04), which was significant in stressed males (p=0.02). We observed a marginally significant trend for a sex x stress interaction (p=0.06) and no effect of sex (p=0.13) (**Figure 2N**).

### 3.2 CUS reduces hypothalamic mitochondrial respiration in a sex-specific manner, but has no effect on mitochondrial complex protein levels, mitochondrial content, or lipid peroxidation

#### 3.2.1 CUS decreases coupled and uncoupled hypothalamic mitochondrial respiration in females but not males

Following euthanasia, we evaluated the effects of stress on mitochondrial function in the hypothalamus as an entry point of the HPA axis. Using high resolution respirometry, we observed significant effects of stress and sex to reduce mitochondrial respiration in the hypothalamus relative to control counterparts (**Figure 3A**). We first observed a main effect of stress to decrease complex I (CI) respiration (p=0.02), which post hoc analysis revealed was driven by stressed females (p=0.04). We did not observe a main effect of sex (p=0.33) nor a sex x stress interaction (p=0.52) in CI respiration. When complex II was activated (CI+CII), these effects persisted, with a main effect of stress (p=0.006), which was significant when comparing CON vs. CUS females only (p=0.01). Again, we saw no main effect of sex (p=0.25) and no interaction (p=.44). We then uncoupled respiration to assess the maximal capacity of the electron transport system (ETS). Here, we observed a main effect of stress to decrease maximal capacity (p=0.02), which post hoc analysis revealed was driven by CUS females (p=0.02). However, we did not observe a main effect of sex (p=0.75) nor sex x stress interaction (p=0.35). Lastly, we inhibited complex I to isolate complex II (ETS CII) activity to determine the contribution of CII to observed respiration reductions. Here, we found main effects of sex (p=0.05) and stress (p=0.04) to decrease complex II respiration, but no sex x stress interaction (p=0.86) (**Figure 3B**), demonstrating effects of stress and sex on both complexes I and II.

**Figure 3.**
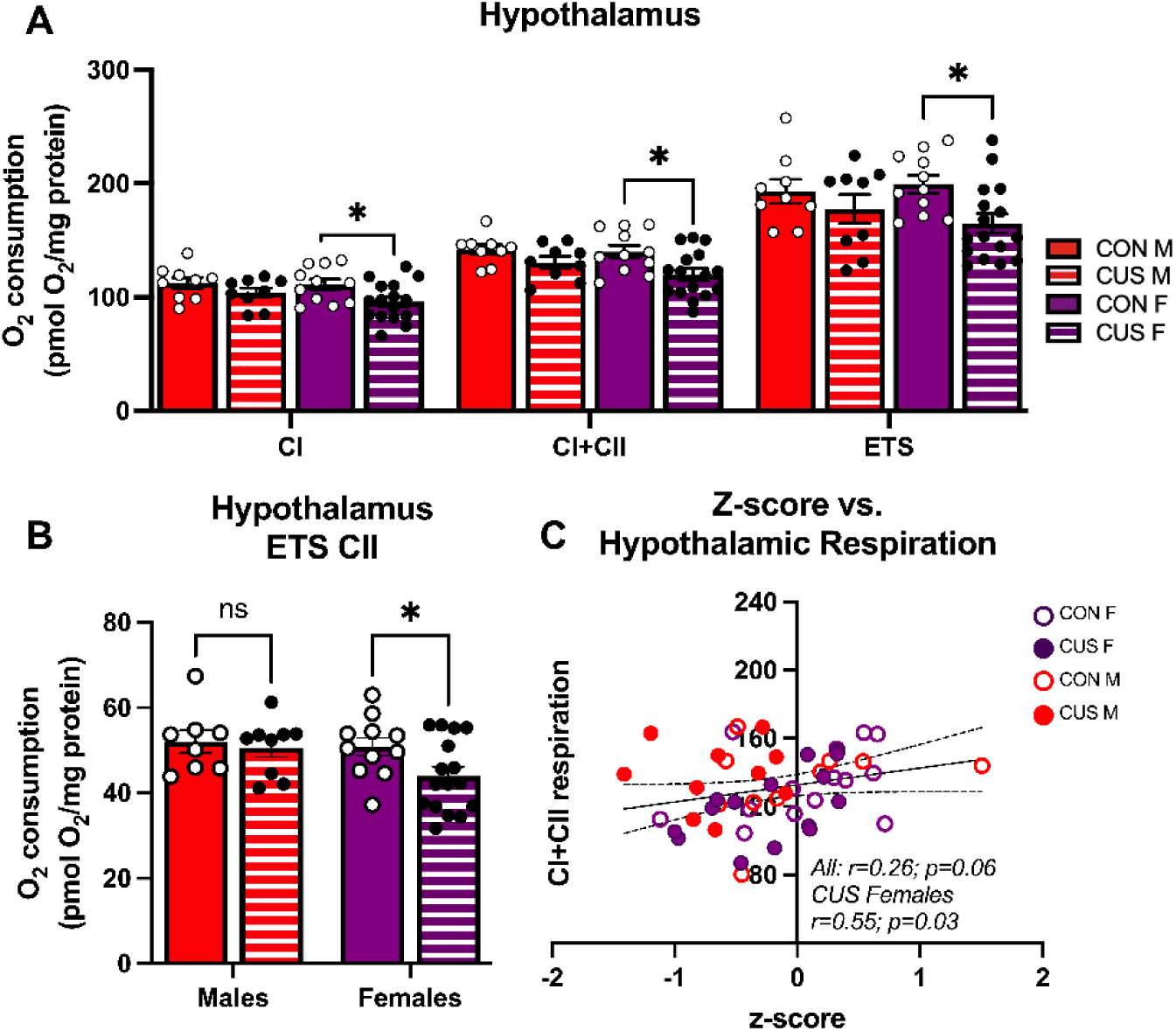
CUS induces sex-specific decreases in mitochondrial respiration in the hypothalamus. **A)** CUS exposure significantly decreased Complex I (CI), Complex I and II (CI+CII), and maximal uncoupled (ETS) respiration in females only. **B)** CUS had no effect on Complex II respiration alone (ETS CII). **C)** There was a significant correlation between hypothalamic CI+CII coupled respiration and the z-score in stressed females only, contributing to the trend observed in all individuals together. n= 12-17/group. Data are represented as ± SEM.

#### 3.2.2 Hypothalamic mitochondrial respiration significantly correlates with composite behavioral z-score in stressed females

We then performed Pearson correlations between coupled respiration of complexes I and II (CI+II) and our computed composite behavioral z-score to determine if there was a relationship between hypothalamic mitochondrial function and behavior. We found a significant correlation between CI+CII respiration and the z-score in stressed females only (p=0.03; r=0.55; **Figure 3C**), such that females with lower respiration also had a lower or more negative z-score. We did not observe this effect with CON females (p=.108, r=.379), with CON males (p=.415, r=.290), nor stressed males (p=.909, r=.041) **(Supplemental Table 3**).

#### 3.2.3 CUS had no effect on mitochondrial complex or content protein expression in males or females

To determine if the effects of CUS were due to differences in mitochondrial OXPHOS complex expression, we analyzed mitochondrial protein expression via western blot (**Figure 4A**). Overall, we did not observe any main effects of stress (CI: p=0.58; CII: p=0.71; CIII: p=0.94; CIV: p=0.37; and CV: p=0.54), effects of sex (CI: p=0.79; CII: p=0.67; CIII: p=0.47; CIV: p=0.55; CV: p=0.94), nor a stress x sex interaction (CI: p=0.79; CII: p=0.74; CIII: p=0.47; CIV: p=0.55; CV: p=0.94) on hypothalamic mitochondrial protein expression (**Figures 4B-F**). Furthermore, we observed no significant differences when we assessed mitochondrial content via TOM20 (stress: p=0.16; sex: p=0.19; interaction: p=0.19) (**Supplemental Figure 2A-B**) and a marker of lipid peroxidation via 4-hydroxynonenal (4HNE) expression (stress: p=0.50; sex: p=0.83; interaction: p=0.85) (**Supplemental Figures 2C-D**).

**Figure 4.**
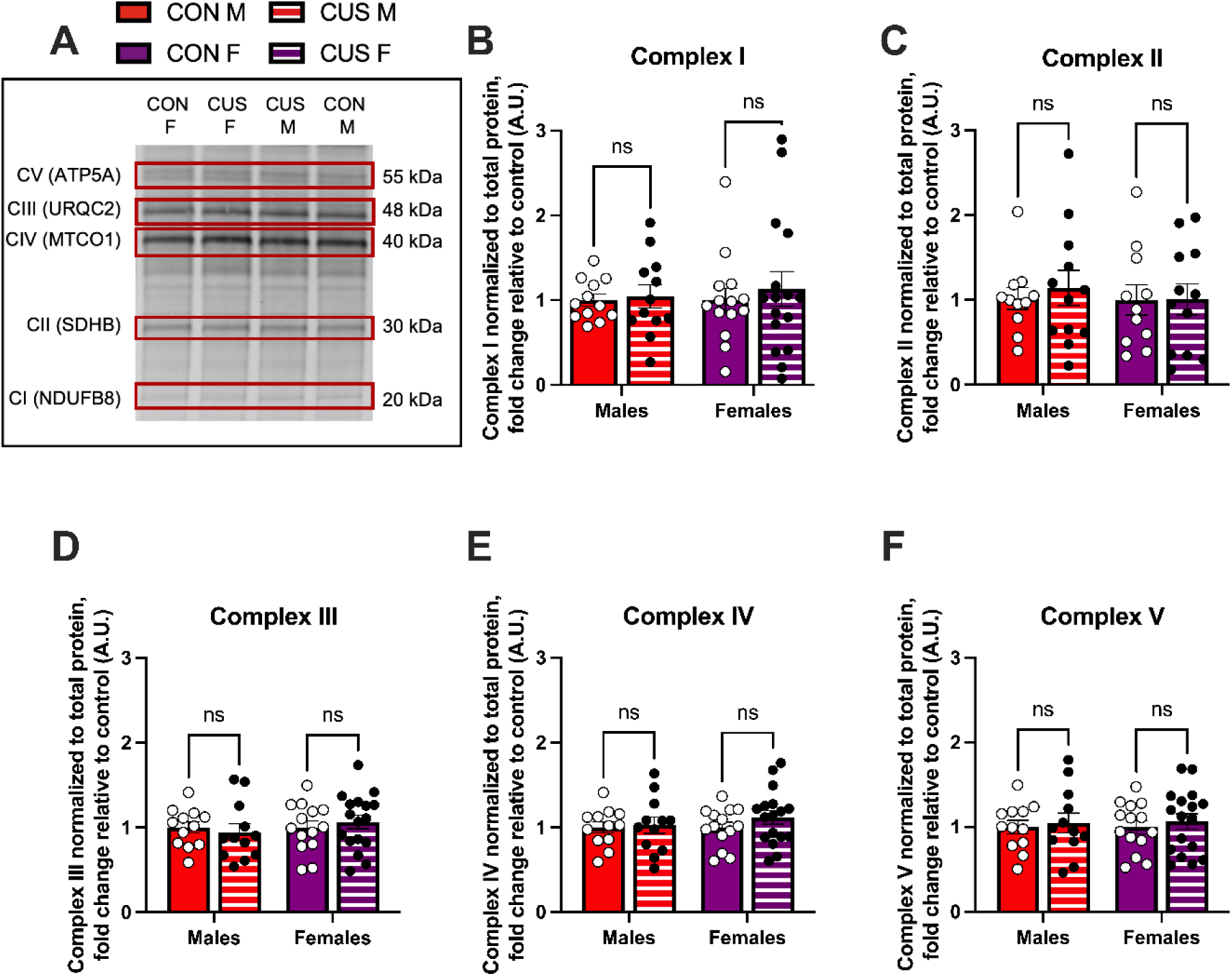
CUS does not alter mitochondrial complex protein levels in the hypothalamus. **A)** Representative western blot image of OXPHOS complex protein expression. There were no significant effects of CUS on hypothalamic mitochondrial **B)** Complex I, **C)** Complex II **D)** Complex III, **E)** Complex IV, and **F)** Complex V protein expression. n= 12-17/group. Data are represented as ± SEM.

### 3.3 CUS reduces adrenal mitochondrial respiration and protein expression, but not mitochondrial content or lipid peroxidation

#### 3.3.1 Females have higher levels of mitochondrial respiration in the adrenal glands

We next assessed the adrenal glands as a region of HPA axis output. When we measured mitochondrial respirometry in the adrenal glands (**Figure 5A**), we observed a main effect of sex (p=0.001) on mitochondrial oxygen consumption coupled to CI. Post hoc analysis revealed significant differences between CON males and females (p=0.01), such that females had higher levels of respiration relative to male counterparts. When complex II was activated, we observed a trend for an effect of sex in the same direction (p=0.08). However, there was no significant effect of sex in uncoupled respiration (p=0.08), suggesting that both sexes have similar adrenal mitochondrial capacities. Notably, when we inhibited complex I to observe complex II activity alone, we found a main effect of sex (p=0.048), where CON females had higher levels of oxygen consumption due to complex II alone compared to males (p=0.03) (**Figure 5B**).

**Figure 5.**
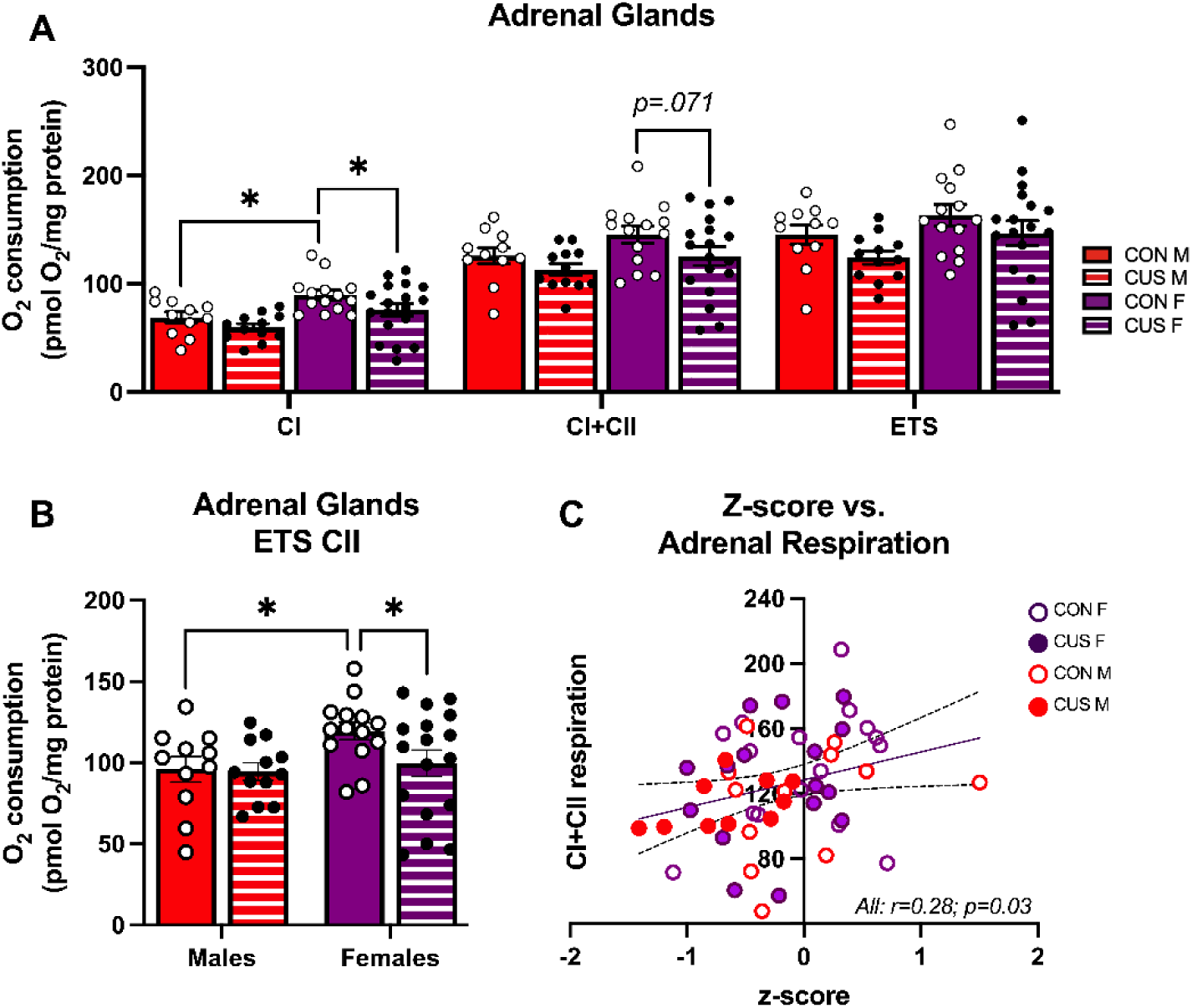
CUS decreases mitochondrial respiration in the adrenal glands. **A)** While sex increased respiration in female controls, CUS exposure significantly decreased Complex I (CI) respiration in females. **B)** When Complex I was inhibited, there was a main effect of sex to increase Complex II respiration (ETS CII), while CUS reduced CII respiration only in females. **C)** There was a significant correlation between adrenal gland CI+CII coupled respiration and the z-score across all individuals. n= 12-18/group. Data are represented as ± SEM

#### 3.3.2 CUS significantly decreases adrenal mitochondrial coupled respiration in females

Our analyses also revealed main effects of CUS. We observed main effects of stress (p=0.03) to decrease CI mitochondrial oxygen consumption, however there was no significant stress x sex interaction (p=0.59) (**Figure 5A**). Post hoc analysis revealed significant differences both between CON and CUS females (p=0.03). We observed a similar main effect of stress in CI+CII coupled respiration (p=0.03), but no stress x sex interaction (p=0.54). Here, post hoc analysis indicated a trend for an effect of stress between CON and CUS females (p=0.07) but not males (p=0.50), suggesting potential sex differences for CII uncoupled respiration. Finally, we observed a marginally significant trend for an effect of stress (p=0.05) on uncoupled respiration, but no interaction (p=0.95), suggesting that maximal capacity may also be somewhat affected by reductions in coupled respiration. When we inhibited complex I to observe complex II activity alone, we found no significant effects of stress (p=0.14). Although the stress x sex interaction was not significant (p=0.20), exploratory post hoc analysis revealed a significant difference between CON and CUS females (p=0.04) (**Figure 5B**).

#### 3.3.3 Adrenal mitochondrial respiration correlates with composite behavioral z-score in stressed males and females

Moreover, when we analyzed adrenal mitochondrial respiration in relation to the z-score, we observed a significant correlation between composite z-score and CI+CII mitochondrial respiration across all individuals (p=0.03; r=0.28) (**Figure 5C**). Notably, this correlation also exists between the z-score and CI mitochondrial respiration (p=0.01, r=0.33). When we separated animals into respective CUS and CON groups, we found the relationship present in CI of CUS animals (p=0.007; r=0.52), but not with CON animals (p=0.38, r=0.17) (**Supplemental Table 3**), suggesting that stressed animals may drive the overall correlation.

#### 3.3.4 CUS increases complex II and III protein expression in males but not females

We examined mitochondrial OXPHOS complex expression in the adrenal glands (**Figures 6A-F**) but did not observe any differences in CI protein expression (stress: p=0.15; sex: p=0.13; and interaction: p=0.63) (**Figure 6B**). We did, however, observe a main effect of stress to increase CII protein expression (p=0.003) (**Figure 6C**), which post hoc analysis revealed to be significant in males (p=0.01), yet no effects of sex (p=0.23) nor an interaction (p=0.23). We also found a main effect of stress to increase CIII protein expression (p=0.01), which was significant in males (p=0.03) (**Figure 6D**), but only a trend for sex (p=0.07) and no significant interaction (p=0.53). There were also no significant effects on CIV protein expression (stress p=0.05; sex: p=0.90; and interaction: p=0.90) (**Figure 6E**) nor CV (stress: p=0.05; sex: p=0.55; and interaction: p=0.55) (**Figure 6F**). Finally, we found no significant effects on TOM20 protein expression (stress: p=0.79; sex: p=0.82; and interaction p=0.82) (**Supplemental Figure 3A-B**) nor 4HNE (stress: p=0.67; sex: p=0.21; and interaction (p=0.21) (**Supplemental Figures 3C-D**), suggesting preserved mitochondrial content and lack of lipid peroxidation.

**Figure 6.**
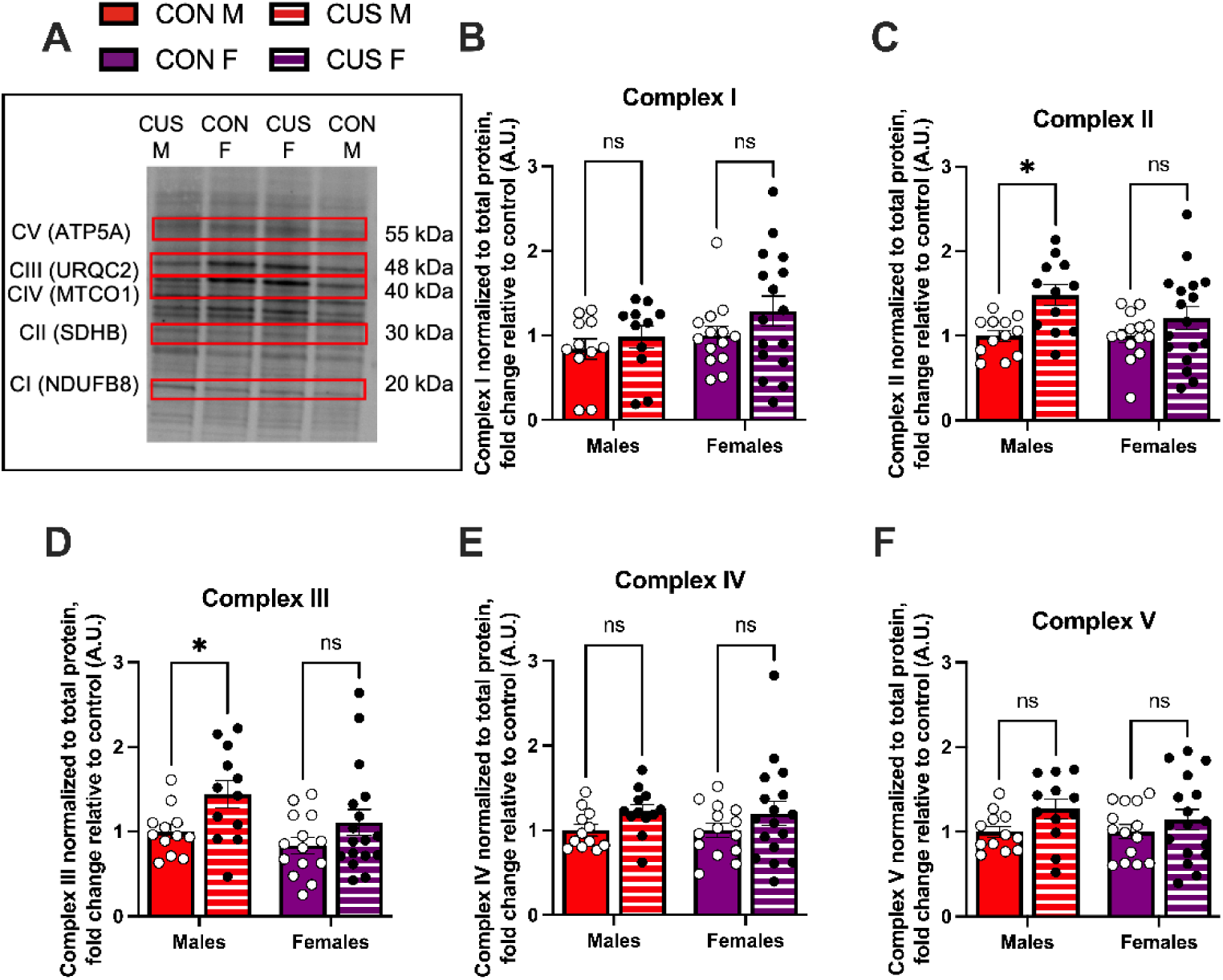
CUS induces sex-specific alterations to mitochondrial complex protein levels. **A)** Representative western blot image of OXPHOS complex protein expression. CUS caused **B)** no impact on Complex I protein expression, **C)** an increase in Complex II protein expression in males, **D)** an increase in Complex III protein expression in males, **E)** no impact on Complex IV protein expression, and **F)** no impact on Complex V protein expression. n= 12-17/group. Data are represented as ± SEM.

## 4. Discussion

Here we show that 28 days of CUS exposure decreased mitochondrial respiration in HPA axis regions and altered adrenal mitochondrial complex protein expression in association with avoidant and passive coping behaviors. Our results were sex- and organ-specific, as we observed effects of CUS in female hypothalamic and adrenal mitochondrial respiration and altered mitochondrial complex protein in male adrenal glands. Notably, we did not observe significant changes in TOM20 or 4HNE, indicating that mitochondrial content and lipid peroxidation were not altered by CUS. This finding suggests that CUS induces functional mitochondrial dysregulation rather than loss of mitochondria or lipid-derived oxidative damage, though involvement of other reactive oxygen species cannot be ruled out. We also observed significant correlations between stress-induced behaviors and mitochondrial respiration in both regions. Our findings indicate a crucial role for functional mitochondria in association with behavioral responses to stress which are dysregulated following chronic stress exposure and highlight sexually dimorphic mechanisms by which stress exerts its effects.

CUS exposure significantly reduced mitochondrial respiration in the hypothalamus in females, without affecting mitochondrial complex protein expression or mitochondrial number. Previous studies described the effects of chronic stress exposure on behavior and brain mitochondrial function in adult male rodents (26–33,42,43), including decreased mitochondrial membrane potential and respiratory rate in mitochondria isolated from the hypothalamus, cortex, and hippocampus (27). These findings have since been corroborated by other studies demonstrating that CUS induces reductions in respiration and membrane potential alongside decreased ATP production (32) and have been linked to pathogenesis of anhedonic and depressive-like behavior (2,7,11). Our work extends the findings of CUS effects on mitochondria to include effects of stress on females. In females, CUS significantly reduced mitochondrial coupled respiration and maximal capacity. As both mitochondrial number and complex protein expression were unaffected, our respiratory findings point to a main effect of stress to disrupt mitochondrial function. Neurons are energy demanding and thus proper mitochondrial function is essential to the survival and performance of these cells (45,46). Hypothalamic mitochondrial function plays a critical role in food consumption, energy expenditure, and adiposity (reviewed in (47)). As chronic stress increases brain energy demands (48) and hypothalamic neuronal excitability (reviewed in (49)), maintaining hypothalamic function becomes critical for proper regulation of energy homeostasis. Decreased mitochondrial function may result in aberrant hypothalamic adaptations to stress, manifesting in behavioral or metabolic pathologies. Female mice did not exhibit the decreased body weight gain that is characteristic of rodent stressors (50) and present in our stressed male mice, suggesting the potential for disrupted hypothalamic regulation of energy expenditure and adiposity. This result agrees with clinical studies demonstrating that women exposed to chronic stress are more vulnerable to metabolic risk than men (51), with stressful life events significantly correlating with increased body mass index in women (52). Additionally, we observed a significant correlation between stress-induced behavior and hypothalamic mitochondrial respiration specifically in stressed females, pointing to a sex-specific relationship between stress-induced behaviors and hypothalamic mitochondrial function. Chronic stress exposure is a risk factor for neuropsychiatric diseases, including depression and anxiety (53), where women are twice as affected (54,55), reporting greater severity of disease and atypical and somatic symptoms (56,57). Thus, our findings suggest that hypothalamic metabolic function may play a role in female vulnerability to stress-induced behavioral pathologies. Our results are in line with recent work by Rosenberg and colleagues (2023) which identified a ‘threat response” network comprised of the hypothalamus and other limbic-associated regions with similar functional mitochondrial signatures that correlated with stress-related behaviors in adult male mice exposed to chronic CORT administration or 10-days of chronic social defeat stress (28). The associations with hypothalamic mitochondrial function and behavior in our study suggest that this threat response network is present and possibly more sensitive in females exposed to chronic stress.

Contrary to prior findings in male rodents (27,32,58) we did not observe disrupted hypothalamic mitochondrial function, or protein expression in stressed males. This discrepancy may reflect differences in CUS protocol design, as our protocol differs from other studies in two key aspects. First, we deliberately excluded any stressors that overtly impacted metabolism to isolate physiological stress effects, including exclusion of food/water restriction, hot/cold stress, and forced swimming. One or more of these metabolic stressors are commonly included in many CUS protocols, including Gong and colleagues (27), who included overnight food and water restriction that may have contributed to the impact on hypothalamic mitochondrial function. Stressors that alter body temperature, physical activity, or caloric intake can trigger alterations in homeostatic regulatory systems, including AMPK signaling within the hypothalamus (reviewed in (59)), inherently impacting mitochondrial processes. Second, the duration of CUS protocol may be another factor. Our CUS protocol was 4 weeks, while others have examined mitochondrial function after 3 (31,43), 4 (32), or 6 (27) weeks. Length of stress exposure has been shown to influence the severity of effects (60,61). Thus, our protocol length and stressor-type appear optimized to detect sex differences in mitochondrial respiratory function that are initiated by physical and psychological non-metabolic stressors. Additionally, we focused on assessing mitochondrial functions specifically within HPA axis regions. It is possible that our CUS protocol impacts male mitochondrial respiratory capacity in higher order regions such as the hippocampus or prefrontal cortex, as these two regions are impacted by CUS protocols (62–64) (reviewed in (65)) and accumulate excess glucocorticoids following chronic stress exposure that decrease function (38). We assessed protein expression limited to mitochondrial complexes (OXPHOS) and content (TOM20) and found no effect of CUS. As CUS alters proteomic pathways related to energy metabolism and glutamate balance (58), it is possible that altered expression in other energy-related proteins compensated and prevented decreases in hypothalamic mitochondrial respiration. Future studies that integrate proteomics with mitochondrial function will be important in characterizing sex differences in the hypothalamic mitochondrial response to chronic stress.

We report here a basal sex difference in adrenal mitochondrial respiration, with females exhibiting higher levels of oxygen consumption than their non-stressed male counterparts. To our knowledge, this is the first report of such a sex difference in mitochondrial function in either rodents or humans. When we analyzed the adrenal glands, we found that CUS caused a significant reduction in mitochondrial respiration, specifically in females. We also observed increased complex II protein expression in males only, and increased complex III protein expression in both males and females. An upregulation of complex II and III proteins may reflect an attempt to increase mitochondrial output within the adrenal glands to compensate for the effects of stress. In males, this strategy may have been effective, as post hoc analyses did not observe significant differences in mitochondrial respiration. However, in females, any compensatory upregulation of complex protein was likely ineffective as we observed a persistent decrease in complex II-dependent respiration following inhibition of complex I. Indeed, a recent study highlights the ability of complex II to activate adrenal complexes III and IV (66), which may explain the lack of effect in males. Complex III protein levels were also upregulated in females, however, a functional deficit in mitochondrial function was still evident in this group. The adrenal glands are enriched with mitochondria, as mitochondria are required for steroid hormone synthesis. Furthermore, the adrenal glands must quickly respond and adapt to the needs of the individual in the face of stress. Thus, a decrease in mitochondrial function could compound the effects of chronic HPA axis activation. Previous work found that cytochrome P450 side chain cleavage (SCC), the rate limiting enzyme for steroid synthesis housed within mitochondria, requires electron chain complexes III and IV to initiate steroidogenesis via complex II activation (66). Thus, an alternative interpretation of the increased complex protein levels could be upregulation to enhance steroidogenesis. However, we did not observe corresponding increased levels of CORT at the same time point. The limited amount of plasma obtained from each mouse prevented us from analyzing additional steroid hormone levels, thus while we believe this discordant relationship between complex protein and mitochondrial function represents a functional mitochondrial deficit, we cannot rule out the possibility of altered steroidogenesis. Adrenal gland mitochondria have previously been shown to respond to stress. Vega-Vasquez et al., (2024) found zone-specific mitotypes across the adrenal cortex based on the steroid hormone they synthesize in adolescent male mice (67). Following chronic psychosocial stress, they found that mitochondrial morphology in the zona glomerulosa and x-zone shifted towards the morphology of those in the zona fasciculata, the region that produces glucocorticoids, while no differences in morphology were observed in the zona fasciculata and no differences in mitochondrial number were observed across layers (67). Our study builds on this literature by providing evidence of functional consequences of chronic stress on adrenal mitochondria across the sexes. Moreover, we identified a significant relationship between adrenal mitochondrial respiration and behavioral phenotype present only in stressed male and female mice, further emphasizing the impact of chronic stress on both male and female mitochondria that may impact behavior. By using a stressor effective in both males and females, we show that adrenal mitochondria are susceptible to the effects of chronic stress, specifically in females, via sex-specific upregulation of complex II protein.

While males exhibited decreased CORT release, females did not exhibit significant effects. One possible explanation could be a floor effect, as female resting CORT levels were significantly lower than that of males, contrasting with previous reports (68). This could stem from the time of collection, as plasma was collected in the morning, when CORT levels in mice are at their lowest. However, the lack of stress effect on female CORT release has previously been reported in the literature. Borrow et al., (2019) assessed morning plasma CORT expression in stressed and non-stressed female C57BL6/J mice and found no differences under basal conditions (40). Thus, it is also possible that female C57BL6/J mice exhibit altered HPA axis responses to stress potentially due to gonadal steroid hormones, specific populations of corticotropin releasing factor (CRF) neurons in key stress-related brain regions, and the distribution of corticosteroid receptors in the brain (69–71). Additionally, it is possible that differences in CORT response would be evident upon acute stress challenge, as reported by others (40). Other measures of HPA axis activation, such as adrenal gland weight, were not significantly affected by stress in either sex, though adrenal gland weights were significantly greater in females compared to males, as reported in (41). The ability of stress to increase adrenal gland weight has mixed support in the literature, with both positive (72) and negative reports (29,30). Together, these data highlight the complexity of sex-specific HPA axis regulation by stress.

The CUS paradigm is a well-validated model for inducing behaviors relevant to anxiety and depression, extensively studied in males (29–33). In females, stress-induced behaviors are more variable and may be strain-dependent (76). Consistent with prior studies (21,39,41,76), we replicated CUS effects in males, including decreased body weight gain, and increased avoidance behavior in the EPM. We also observed significant sex-specific effects of stress-induced behavior where stressed females, in addition to EPM avoidance behavior, also exhibited passive-coping behavior in the FST, but did not exhibit reduced body weight gain. Several studies have similarly reported similar responses either in weight gain (41) or avoidance behavior (77). As previous reports have suggested that age of assessment impacts findings (77), it is possible that CUS may enhance anxiogenic behaviors in older females. Neither group showed deficits in the OFT, though this could be attributed to differences in testing conditions as we use low lighting conditions (8-10 lux) to assess inherent avoidance behavior in lieu of induced avoidance behavior. We incorporated these behavioral measures into a z-score to assess the presence of stress-induced endophenotypes. Previous studies have shown that stress provokes behavioral phenotypes in males (34–36), though few have assessed this parameter in females (35). Here, we show that CUS induces a robust composite behavioral phenotype in males, as reflected in a decreased behavioral z-score. In females, while the composite z-score was not significantly altered, CUS did elicit increased passive coping in the FST. This highlights the importance of evaluating stress-induced phenotypes using both individual tests and integrative measures, as females may exhibit domain-specific alterations that are not fully captured by composite z-score approaches. Our behavioral testing was limited to avoid interference with downstream analyses. Thus, it is possible that our CUS protocol induces significant behavioral alterations in females that occur in other behavioral domains (*e.g.,* cognition, sociability) not explored here. Interestingly, while CUS did not impact composite z-score in females, we still observed significant correlations between the z-score and hypothalamic mitochondrial respiration specifically in stressed females. Similarly, we observed significant correlations between adrenal gland mitochondrial respiration and z-score that was driven particularly by stressed groups. Together, these associations suggest that mitochondrial function is regionally specialized and stress-responsive and CUS reorganizes regional mitochondrial function to predict behavioral outcomes.

### 4.1 Limitations

Our well-controlled experimental design provides valuable insight into how chronic stress alters mitochondrial function across the HPA axis, yet several potential limitations offer important directions for future research. First, plasma and tissue regions were limited as we prioritized individual responses over pooled samples. Thus, comprehensive analyses of steroid hormones and analysis of mouse pituitary gland, the remaining key node of the HPA axis, were not possible, but remain a future objective. Second, the hypothalamus has heterogeneous cell populations with divergent physiological functions (e.g., orexigenic neuropeptides such as Agouti-related protein (AgRP) and Neuropeptide Y (NPY) versus anorexigenic neuropeptides cocaine- and amphetamine-regulated transcript (CART) and α-melanocyte-stimulating hormone (α-MSH, a product of proopiomelanocortin (POMC) processing) (78) that are affected by stress (49). Our study investigated whole hypothalamic homogenates. This approach enabled broad regional profiling but precluded cell-type specific populations. Previous work identified that 10 days of CUS decreased AgRP neuronal function in both males and females (61). As mitochondrial function provides energy to sustain neuronal firing (46), future studies should evaluate whether stress-induced mitochondrial alterations are confined to specific hypothalamic cell populations. Finally, although we identified robust sex- and organ-specific effects of CUS on mitochondrial respiration, we did not assess the molecular pathways or signaling mechanisms driving these effects. Stress-related changes in mitochondrial function may be mediated by glucocorticoid receptor activation, oxidative stress, neuroinflammation, or neurotrophic signaling (e.g., BDNF), which may differ between males and females. Investigating these upstream regulators will be critical for elucidating mechanisms underlying stress sensitivity and resilience, potentially identifying additional targets for therapeutic intervention.

## 5. Conclusions

Our findings demonstrate that CUS leads to sex-specific disruptions in mitochondrial function within key regions of the HPA axis. Specifically, stress exposure significantly impaired hypothalamic and adrenal mitochondrial respiration in females, accompanied by region- and sex-dependent alterations in mitochondrial complex protein expression. Importantly, these mitochondrial changes were significantly associated with stress-related behavioral outcomes, suggesting a mechanistic link between energy metabolism and behavioral adaptation to stress. Our data underscore the critical role of mitochondria in mediating the effects of chronic stress and highlight sex-dependent differences in metabolic vulnerability. Clinically, our findings suggest that mitochondrial function may represent a novel biomarker or therapeutic target for stress-related neuropsychiatric and/or metabolic disorders, particularly in women, who are disproportionately affected by chronic stress. Understanding sex-specific mitochondrial adaptations to chronic stress may help guide the development of personalized interventions aimed at mitigating stress-induced dysfunction across central and peripheral systems. Overall, our results support the existence of distinct mitochondrial mechanisms through which chronic stress differentially impacts males and females, potentially contributing to sex-specific trajectories of stress-related disorders.

## Data Availability

All data and protocols presented in the manuscript are available upon request.

## CRediT Author Contributions

AMC: Conceptualization, data curation, formal analysis, investigation, methodology, project administration, supervision, validation, visualization, writing—original draft, writing— review and editing. NIF: Conceptualization, data curation, formal analysis, investigation, methodology, project administration, writing – review and editing; AM: data curation, investigation, writing – review and editing; AC: data curation, investigation, writing – review and editing; RRDP: data curation, investigation, writing – review and editing; CF: data curation, investigation, writing – review and editing; LF: data curation, investigation, supervision, project administration, writing – review and editing; EC: data curation, investigation, writing – review and editing; JG: data curation, investigation, methodology; writing – review and editing; TLP: data curation, investigation, writing – review and editing; SW: data curation, investigation, writing – review and editing; FP: data curation, investigation, project administration, supervision, writing – review and editing; FGS, RCW, SKW, MJR: Conceptualization, formal analysis, funding acquisition, methodology, project administration, resources, software, supervision, writing— review and editing; FH: Conceptualization, data curation, formal analysis, funding acquisition, investigation, methodology, project administration, resources, software, supervision, validation, visualization, writing—original draft, writing— review and editing.

## Acknowledgments

The authors would like to thank Drs. Lawrence Reagan and Raoni Dos Santos for their helpful discussions and feedback. The authors gratefully acknowledge the USC DLAR staff for their care and attention to our animals and equipment. Some figures were made using BioRender.com.

