## Supplementary Figure 1 for "Sex-specific effects of chronic unpredictable stress on mitochondrial function in the HPA axis in mice"

University of South Carolina School of Medicine

Columbia VA Health Care System

6439 Garners Ferry Road

Columbia, SC 29209

**Supplemental Figures**

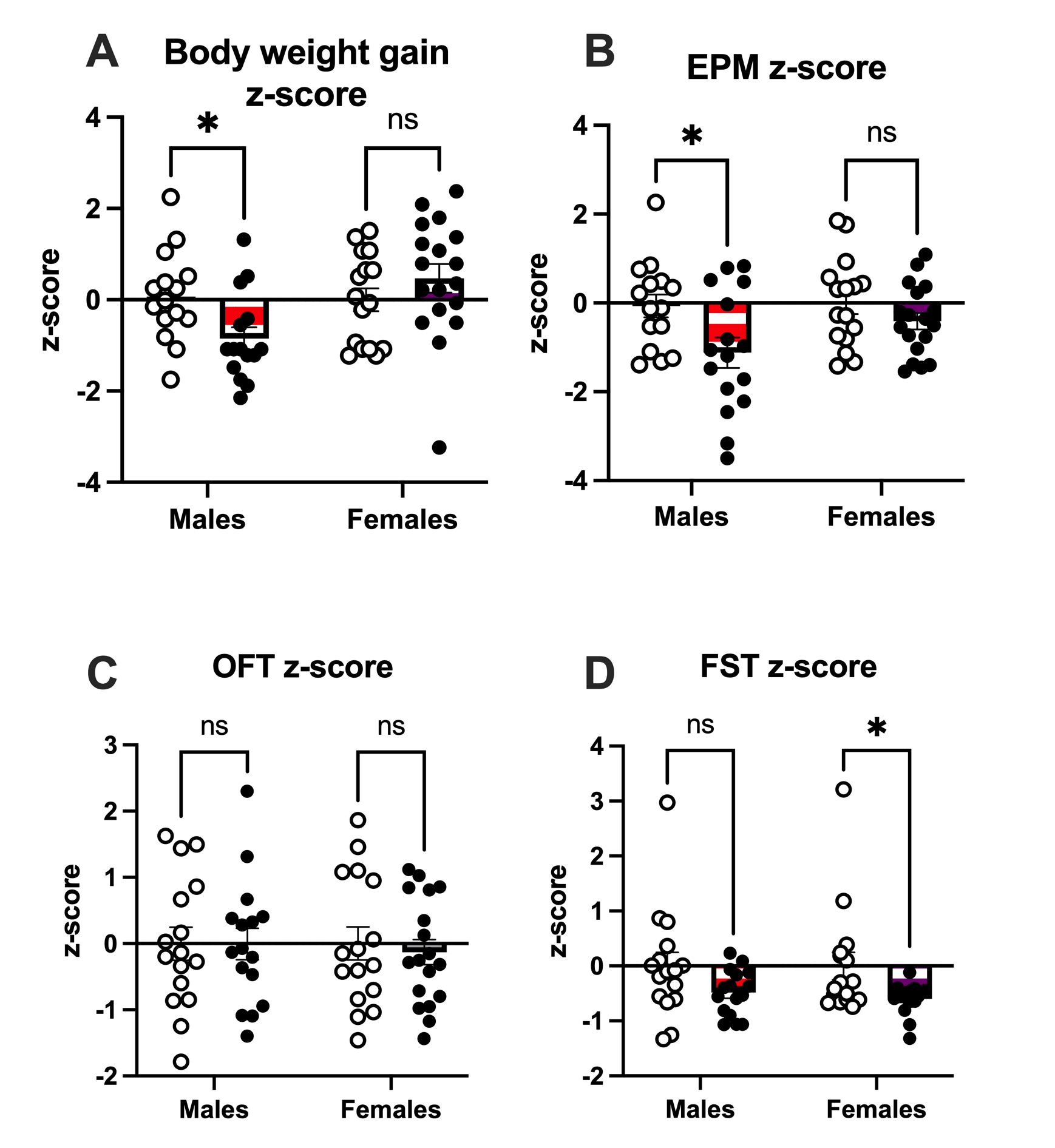

**Supplemental Figure 1. Sex-specific differences in individual test z-scores based on behavioral task. A)** CUS males had a significantly negative z-score compared to controls, but there were no differences between females. **B)** In the EPM, CUS males had a significantly negative z-score compared to controls, but females were not affected. **C)** There was no effect of CUS on z-score in the OFT. **D)** In the FST, CUS females had a significantly negative z-score compared to controls, but males were not affected. n= 14-18/group. Data are represented as ± SEM.

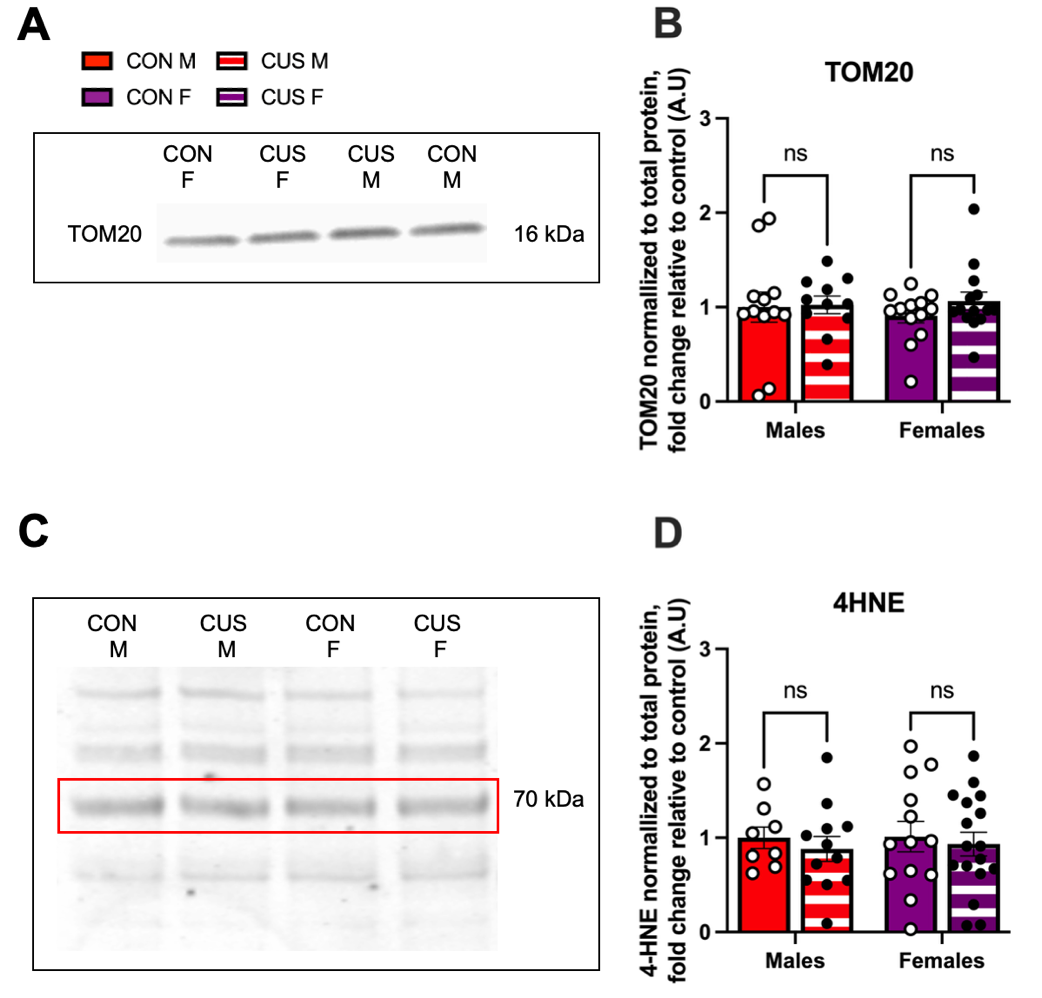

**Supplemental Figure 2. CUS does not alter mitochondrial content or lipid peroxidation in the hypothalamus. A)** Representative image of TOM20 protein expression. **B)** CUS did not alter mitochondrial content via TOM20 protein expression. **C)** Representative image of 4-hydroxynonenal (4HNE) protein expression. **D)** CUS did not alter 4HNE protein expression. n= 12-17/group. Data are represented as ± SEM.

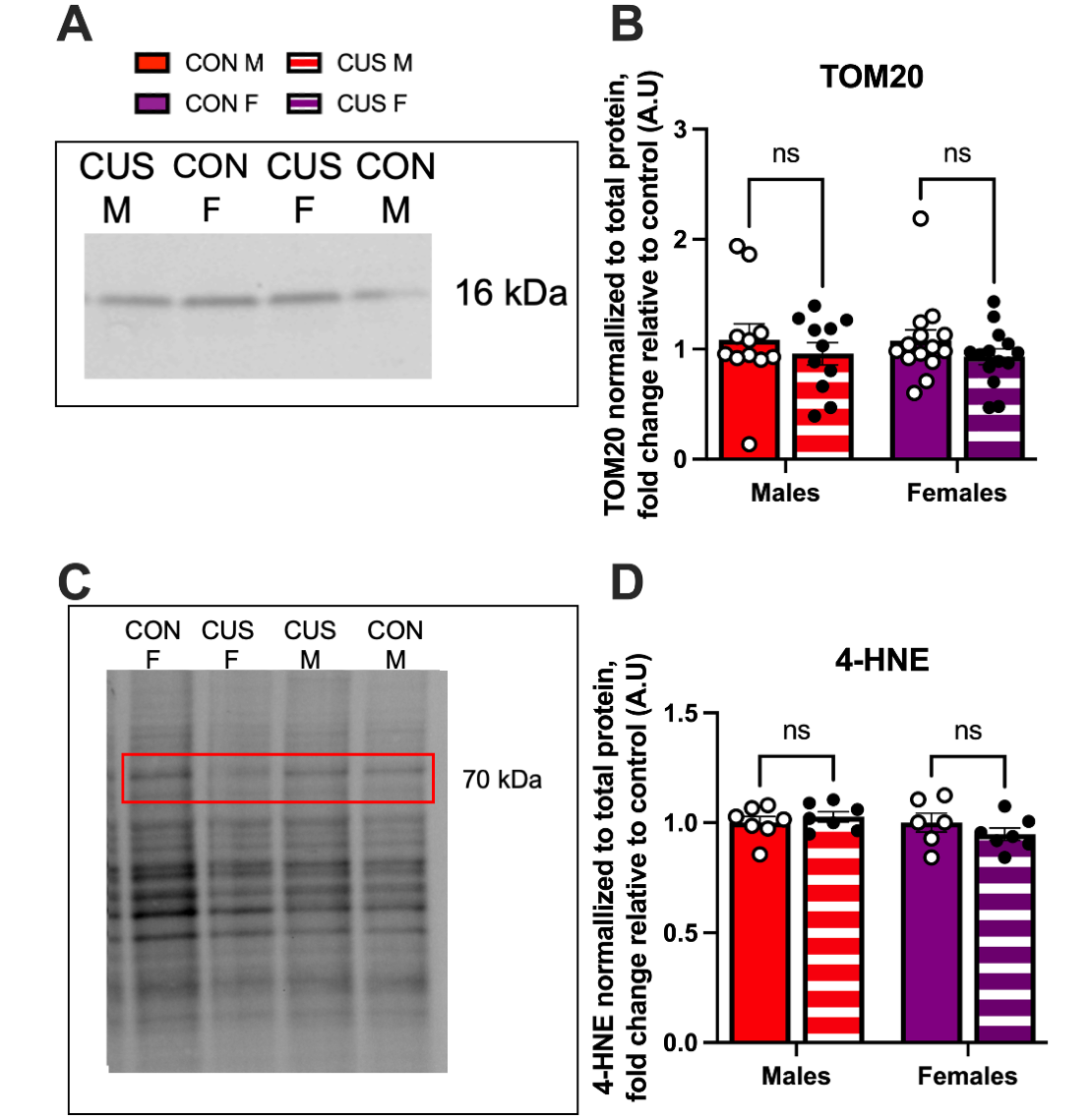

**Supplemental Figure 3. CUS does not alter mitochondrial content or lipid peroxidation in the adrenal glands. A)** Representative image of TOM20 protein expression. **B)** CUS did not alter mitochondrial content via TOM20 protein expression. **C)** Representative image of 4HNE protein expression. **D)** CUS did not alter 4-hydroxynonenal (4HNE) protein expression. n= 6-17/group. Data are represented as ± SEM.

**Supplemental Table 1.** Statistical details for main text analyses.

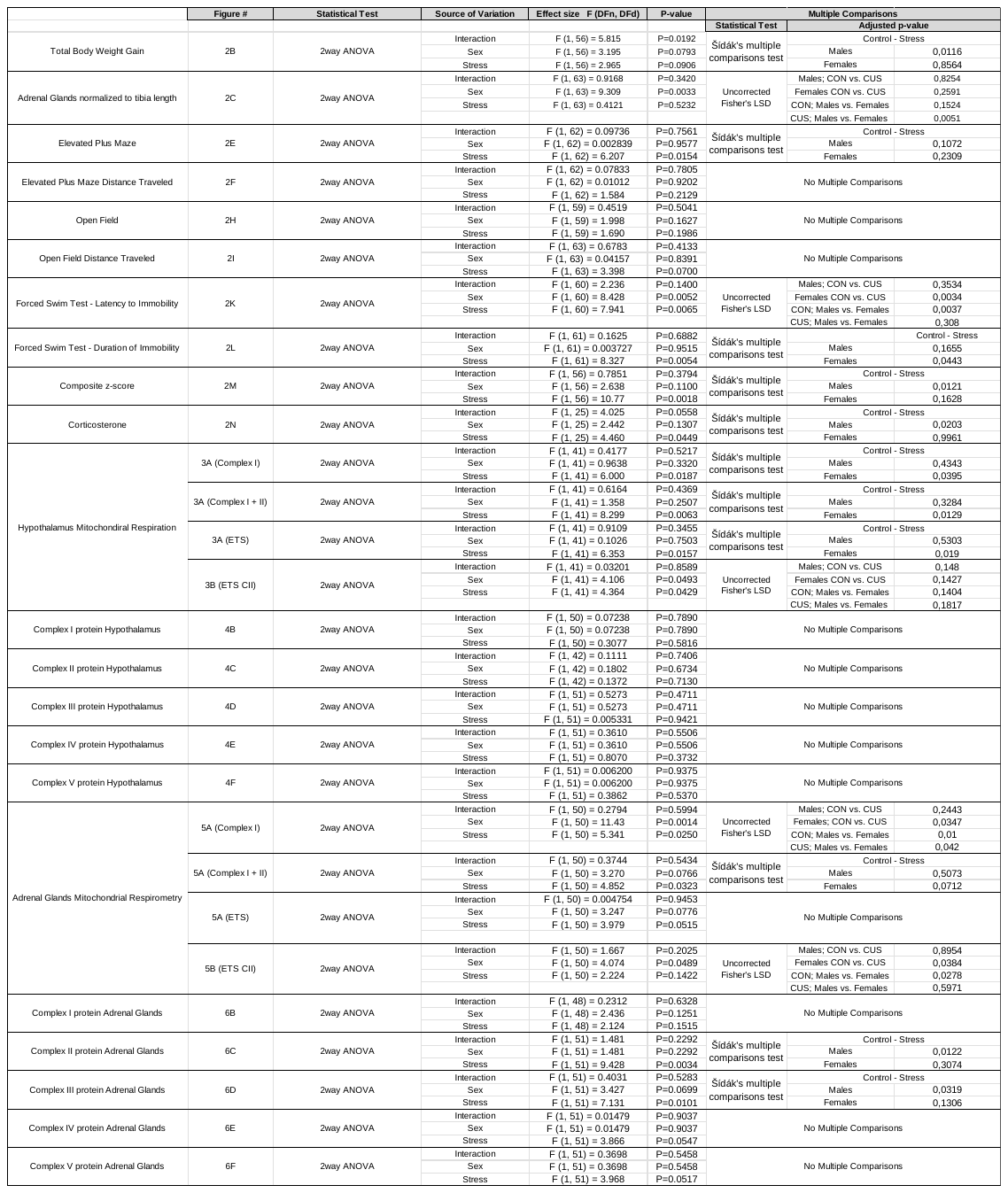

**Supplemental Table 2.** Statistical details for supplemental figure analyses.

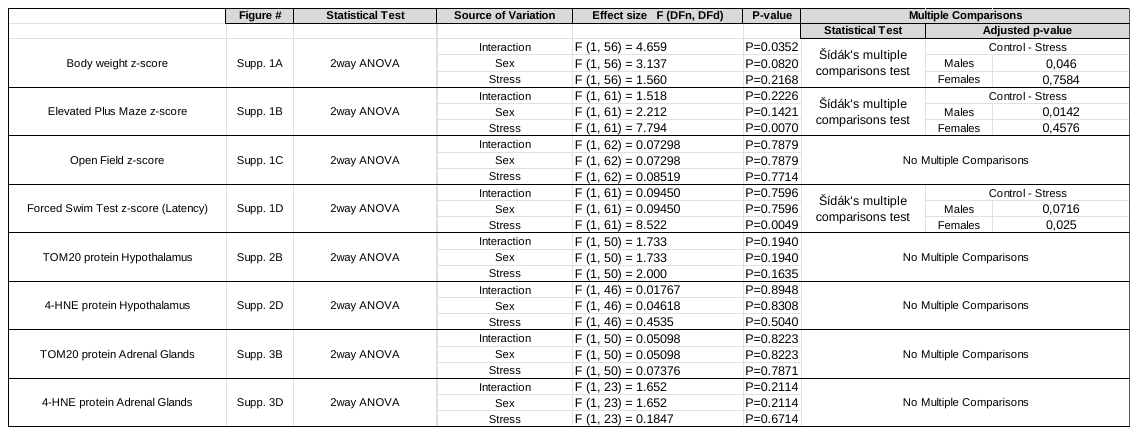

**Supplemental Table 3.** Correlations between composite behavioral z-score and mitochondrial respiration across individual groups and CON vs. CUS groups in the hypothalamus and adrenal glands.

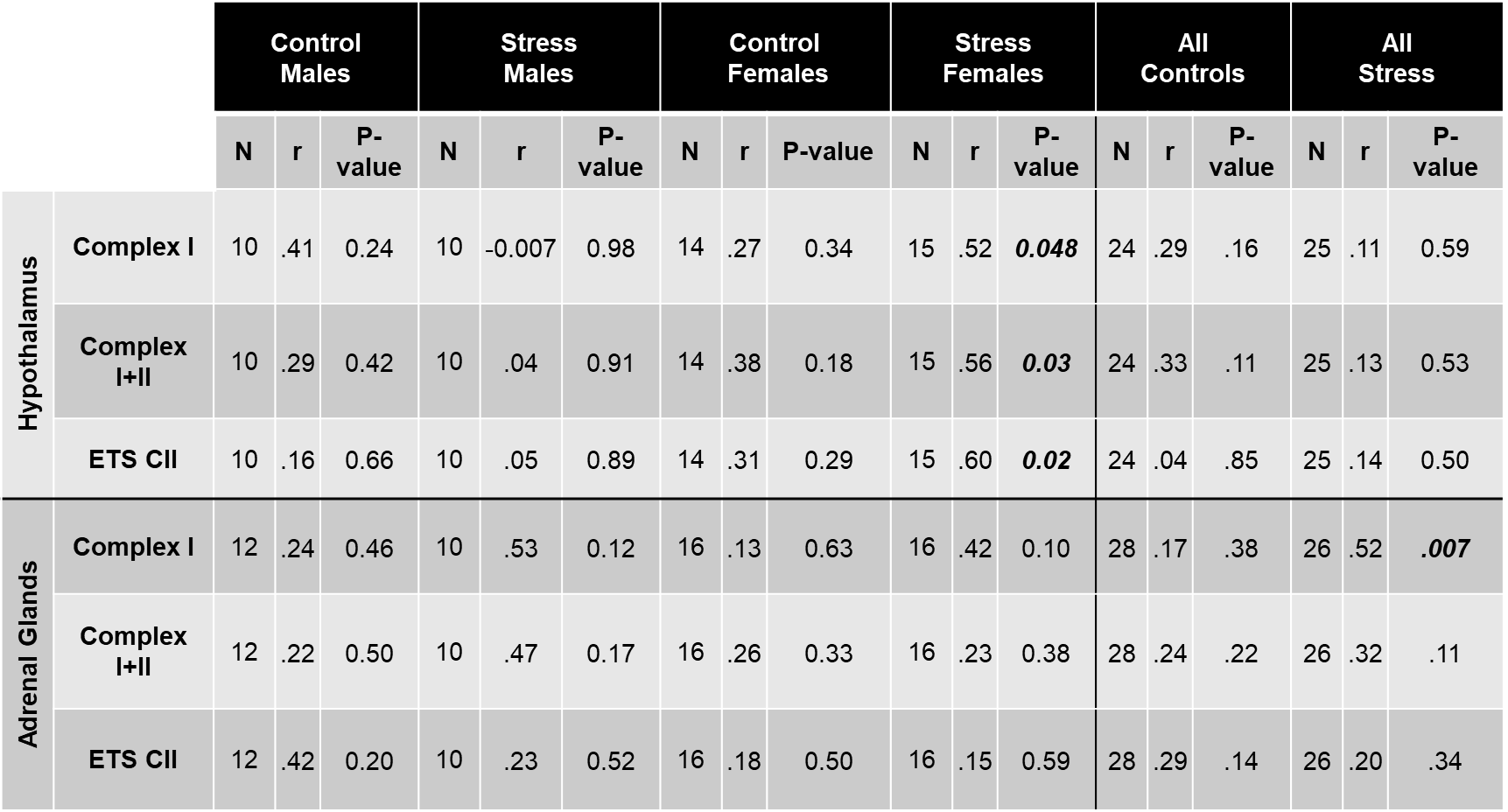
